# High-throughput microbiome profiling and co-occurrence with antibiotic resistance genes in *Lucilia sericata*

**DOI:** 10.1101/2025.06.11.659062

**Authors:** Arwa Shatta, Xavier Chavarria, Jun Ho Choi, Singeun Oh, Myungjun Kim, Dongjun Kang, Yoon Hee Cho, Du-Yeol Choi, Myung-hee Yi, Ju Yeong Kim

## Abstract

Blow flies such as *Lucilia sericata* (Diptera: Calliphoridae) serve important ecological functions as decomposers. However, due to their close association with decaying organic matter they also play potential roles as reservoirs for pathogenic bacteria and antimicrobial resistance genes (ARGs). In this study, we characterized the bacterial communities and resistome profiles of *L. sericata* specimens collected from six provinces across South Korea using 16S rRNA gene metabarcoding and targeted PCR screening. The microbiome was dominated by *Dysgonomonas, Vagococcus, Pseudomonas, Ignatzschineria*, and *Providencia* with geographic variation in community structure. Flies from Chungnam exhibited the lowest microbial diversity, while samples from Jeonnam and Gyeonggi showed greater richness and evenness. Beta diversity analyses confirmed geographic structuring of bacterial communities, with semi-urban, rural locations harboring more diverse taxa. Notably, opportunistic pathogens such as *Proteus mirabilis* and *Providencia* were detected, alongside a range of ARGs (*blaTEM*, *ermB*, *sul1*, *aac(6′)-Ib-cr*, *cat* and *mecA)* and integron elements (*intI* and *intII*), suggesting that *L. sericata* may act as a reservoir of clinically important microbes and resistance genes.

**Importance:** The environment plays a significant role in shaping the microbiome of flies. Due to their motile nature and close association with decomposing matter blow flies of the species *Lucilia sericata* can harbor diverse bacterial communities, including potential pathogens that threaten human and animal health. Furthermore, due to their synanthropic lifestyle, this species is also exposed to bacteria that host antibiotic resistance genes (ARGs). Here we investigate the role of blowflies as reservoirs of potential pathogens and ARGs using metabarcoding. Our study revealed a diverse microbiome and resistome shaped by location and possible biotic or abiotic factors. These findings provide baseline information for wildlife surveillance and emphasize the importance of including synanthropic flies in strategies aimed at controlling the dissemination of antibiotic resistance.

## Introduction

Green bottle flies are saprophagous blow flies commonly found in environments rich in decomposing organic material (1). They are relatively large calyptrate flies, recognized by their metallic blue or green color and are widely distributed across human-associated environments (2). Blow flies of the species *Lucilia sericata* are known for their ecological role as decomposers and their close and frequent contact with decaying organic matter (3), which make them potential mechanical vectors of pathogens (4) and reservoirs of antimicrobial resistance genes (ARGs) (5). Previous studies have reported the bacterial community and the presence of antimicrobial resistance genes (ARGs) in the microbiota of house flies (*Musca domestica*) (4, 6, 7), whereas relatively little research has focused on the occurrence and distribution of ARGs in other flies. *L. sericata* is of a particular interest due to its widespread distribution, synanthropic behavior, and importance in medical (8) and forensic entomology (9). While they have been traditionally studied for their therapeutic use (10). Studies suggest that blow flies commonly harbor pathogenic bacteria associated with ARGs within its gut and on their exoskeleton acquired from contaminated environments (3, 11). This raises the possibility of *L. sericata* serving as unrecognized vectors of resistant bacteria, especially in urban and agricultural settings.

Since *L. sericata* commonly colonize decaying matter such as carcasses and excrement, they likely acquire a sizeable proportion of their microbiome from these sources and then serve as mechanical vectors transferring them to other hosts including humans (3). These flies harbor a rich commensal microbiome encompassing species such as *Providencia*, *Ignatzschineria*, *Vagococcus*, and *Myroides,* some of which have been shown to be transmitted horizontally (11). Commensals such as *Proteus mirabilis* aid larval digestion and suppress pathogenic microbes by secreting antimicrobial agents (12). Similarly, *Providencia* are suggested to contribute to maggot development through the biosynthesis of the essential amino acid methionine (13). Notably, both genera are also known opportunistic human pathogens (14), highlight the dual role of *L. sericata* as beneficial organisms in specific contexts and potential public health risks.

In this study, we employed 16S rRNA gene amplicon Illumina sequencing to characterize the bacterial communities present in blow flies of the species *L. sericata* collected from garbage waste in across South Korea. Additional PCR-based screening was conducted to detect the presence of the most common antimicrobial resistance genes (ARGs), to assess their prevalence in these flies. This study aimed to explore the influence of different location in the diversity and composition of bacterial communities in *L. sericata,* along with the prevalence of potential pathogenic taxa and antimicrobial resistance genes.

## Methods

### Blowflies Sampling, DNA extraction and molecular identification

A total of 129 flies were collected from various environments across six regions in South Korea between April 2023 and June 2023 (Supplementary Figure S1). Sampling locations were selected based on their potential to attract flies, particularly areas with organic waste accumulation (supplementary Table S1). Flies were captured using sweep nets, immediately transferred into sterile containers, and transported to the laboratory in RNAlater (Invitrogen, Vilnius, Lithuania) for further processing. Samples were stored at −80 °C. Prior to DNA extraction, flies were washed with absolute ethanol to remove potential environmental contamination. Genomic DNA was extracted from whole-body homogenates of individual blowflies using a NucleoSpin DNA Insect Extraction Kit (Macherey-Nagel, Düren, Germany) following the manufacturer’s instructions. DNA was then stored at −20 °C. Species identification was conducted through morphological assessment and confirmed by Sanger sequencing (Bionics, Seoul, Republic of Korea) of the mitochondrial cytochrome c oxidase subunit I (MT-CO1) partial sequence. Sequencing was performed using LCO1490/ HCO2198 primer pair, following the protocol described by Yi. et al (15).

### High-throughput sequencing of 16S rRNA gene amplicons

The bacterial microbiome was identified through metabarcoding of the bacterial 16S rRNA V4 region amplified using the 16S_V4_515F (5′-GTG CCA GCM GCC GCG GTA A-3′) and 16S_V4_806R (5′-GGA CTA CHV GGG TWT CTA AT-3′) primers, generating a 300 bp amplicon. Forward (5′-TCG TCG GCA GCG TCA GAT GTG TAT AAG AGA CAG-3′) and reverse (5′-GTC TCG TGG GCT CGG AGA TGT GTA TAA GAG ACA G-3′) Illumina adapters were synthethized upstream of the corresponding primer for Illumina compatibility. PCR amplification was performed using KAPA HiFi HotStart ReadyMix (Roche Sequencing Solutions, Pleasanton, CA, USA) under the following conditions: initial denaturation at 95 °C for 5 min, 25 cycles at 98 °C for 30 s, 55 °C for 30 s, 72 °C for 30 s, and final extension at 72 °C for 5 min.

After PCR, AMPure XP (Beckman Coulter, Brea, CA, USA) was used for amplicon purification. A short-cycle (8 cycles) amplification step was performed to attach Illumina multiplexing indices. DNA concentration for each sample was measured using the QuantiFluor ONE dsDNA System (Promega, Madison, WI, USA), and amplicons were pooled in Tris-HCl buffer to create a 2 nM pool, subsequently diluted to a final concentration of 50 pM. The pooled library was sequenced in the Illumina iSeq100 sequencing system using the Illumina iSeq™ 100 i1 Reagent v2 kit (Illumina Inc., San Diego, CA, USA) following the manufacturer’s protocol. PhiX Control v3 (Illumina) was used as for quality control and calibration of the sequencing runs.

### Bioinformatics and statistics

Posterior bioinformatic analyses were conducted using the QIIME 2 pipeline v. 2022.11.1 (https://qiime2.org/). Raw reads were demultiplexed and trimmed with q2-cutadapt and then denoised using DADA2 using the consensus method for chimera exclusion. Taxonomic classification of the amplicon sequence variants (ASVs) was performed using a classify-consensus-blast classifier against the SILVA v. 138.99 database. Reads were rarefied to 10,000 reads for the subsequent calculation of alpha and beta diversity metrics. Statistical comparisons of alpha diversity among fly populations were conducted using the Kruskal–Wallis rank sum test, as implemented in the microeco package (16) in the R software v. 4.3.1 (https://www.R-project.org/). Post hoc pairwise differences were assessed using pairwise Wilcoxon rank-sum tests with Benjamini–Hochberg false discovery rate correction.

Beta diversity differences within and between the fly specimens were assessed via permutational multivariate analysis of variance (PERMANOVA) using QIIME 2. Ordination of beta diversity distances was performed through Principal Coordinate Analysis (PCoA) in microeco. To identify representative taxa per region, differential abundance was calculated with the Linear Discriminant Analysis (LDA) Effect Size (LEfSe) conducted in R using the microeco trans_diff class.

### Bacterial specific identification and antibiotic resistance gene screening

The presence of *Proteus mirabilis* and *Providencia* spp., two opportunistic pathogens commonly found together in flies (3, 17), was tested through specific PCR analyses. Primers for the *ureR* gene were used to detect *P. mirabilis* at the species level, while the genus *Providencia* was detected at the using partial gene sequence of the16S rRNA gene (Table 1). Contigs of the *Providencia* amplicons were obtained in Geneious Prime v.2025.1.1, and matrices of the contigs and sequences of the other species of the genera obtained from NCBI were aligned in Clustal X v.2.1. The alignments were cut to equal length and translated to proteins to check for indels and stop codons in Mesquite v.3.81. The resulting file was used to construct a Bayesian tree of the *Providencia* genus showing the position of the *Providencia* amplicons obtained through Sanger sequencing. Markov Chain Monte Carlo (MCMC) analysis was run for 100 million tree generations with sampling set every 10,000 trees with 10% burnin in BEAST X v.10.5.0. Maximum likelihood supports were calculated in the IQTREE web server using 10,000 ultrafast bootstrap analysis and 1000 maxit iterations. Substitution model selection was performed in IQTREE v.3.0.1. The resulting tree was visualized and edited in FigTree v.1.4.4.

**Table 1.**
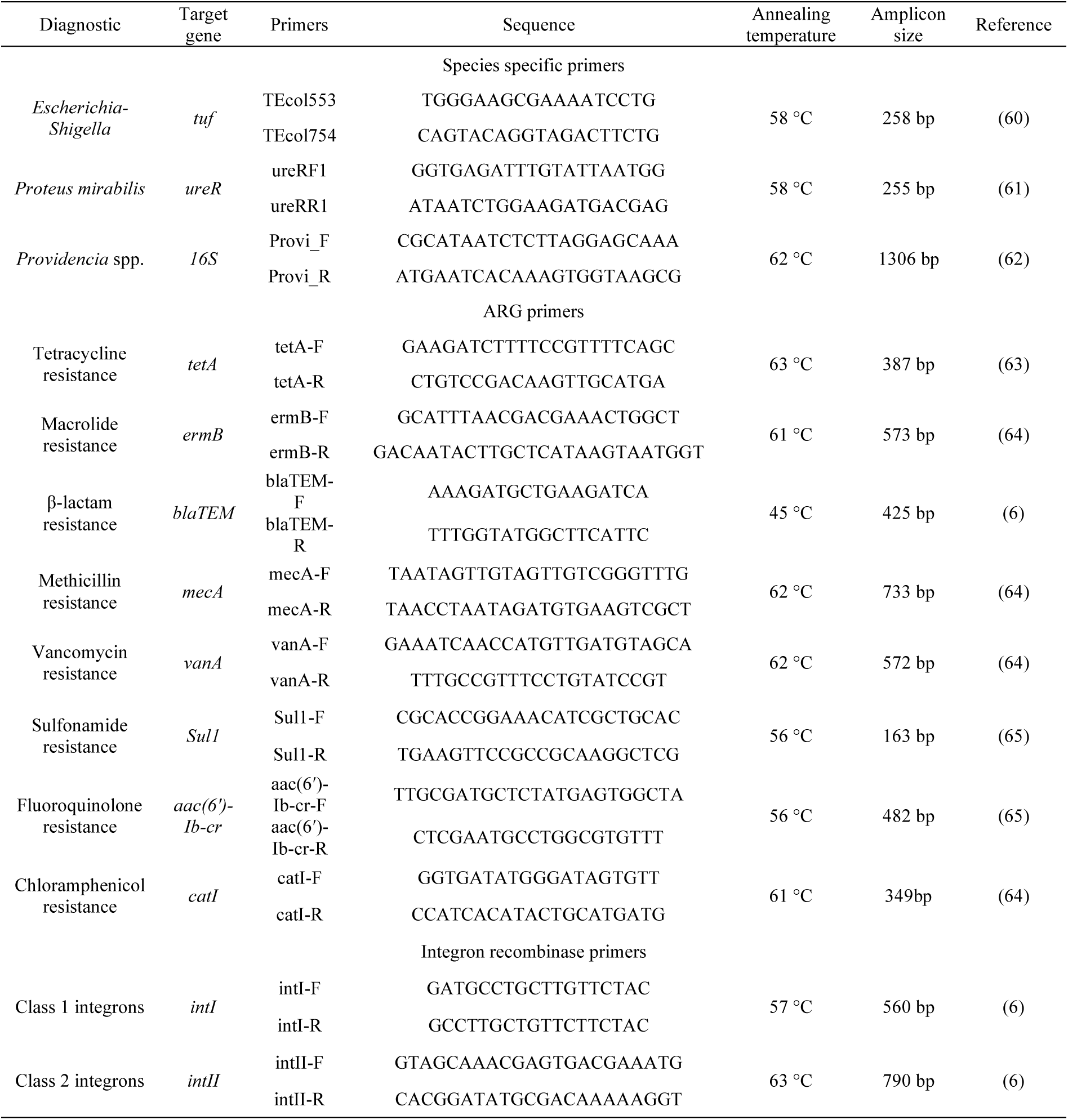
PCR primer sequences used for specific gene detection in this study.

**Table 2.**
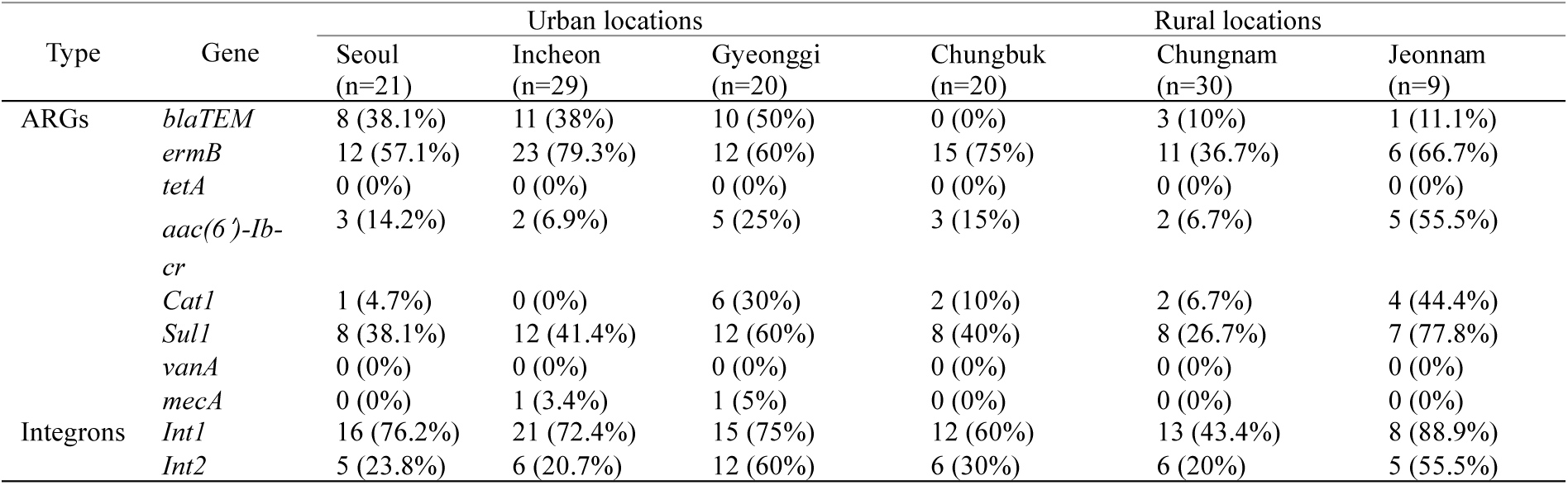
Antibiotic resistance genes and integrons detected in *Lucilia sericata* from six provinces in South Korea.

The presence of common antibiotic resistance genes in the microbiome of flies was assessed using gene-specific PCR assays. Eight different resistance genes were tested, each corresponding to a important antibiotic classes. Tetracycline resistance was evaluated through the *tetA* gene, while macrolide resistance was assessed with the *ermB* gene. For β-lactam resistance, the *blaTEM* gene was tested, while methicillin resistance was determined through the *mecA* gene. Vancomycin resistance was assessed using the *vanA* gene. Sulfonamide resistance was evaluated by testing for the *sul1* gene, while aminoglycoside resistance was determined through the *aac(6′)-Ib-cr* gene. Finally, chloramphenicol resistance was tested using *cat1* gene primers. Furthermore, since integrons are key genetic elements that facilitate the integration and horizontal transmission of certain gene cassettes, including those conferring resistance to multiple antibiotics (18), they were also screened alongside the antibiotic resistance genes. The presence of integrons sequences was screened through PCR of the *intI* recombinase gene for class 1 integrons and the *intII* recombinase gene for class 2 integrons. The presence and absence of ARGs and integrons were compared by location through pairwise PERMANOVA based on Jaccard distances using the R package vegan.

PCR screenings were performed using AccuPower PCR PreMix (Bioneer, South Korea) with each primer at a final concentration of 0.5 µM in a total reaction volume of 20 µL. Amplification conditions were: an initial denaturation at 95°C for 6 min, 35 cycles of 95°C for 30 s, primer-specific annealing for 30 s, and 72°C for 30 s, with a final extension at 72°C for 6 min. PCR grade water was used instead of DNA for negative controls. Amplicons were visualized in 1% gel agarose electrophoresis. Amplicons with bands of the expected target size were purified using the same method as the metabarcoding cleanup step and sequenced to confirm ARG identity (Bionics, South Korea). Primer sequences and their annealing temperatures used for these assays are shown in Table 1.

### Microbiome and antibiotic resistance gene correlation network

The patterns of bacterial ASVs and ARG co-occurrence were assessed using a Spearman correlation network with the Hmisc package in R. The top 100 most abundant ASVs were selected, normalized to relative abundances and then centered log-ratio (CLR) transformed with a pseudocount of 1e-6 to account for composition constraints. Correlations between ARGs and ASVs with an absolute coefficient > 0.2 and a p-value < 0.01 were considered significant. Network visualization of the significant correlations was generated using the igraph package.

## Results

### Bacterial microbiome composition

A total of 3,025,117 reads were obtained with an average count of 23,450 ± 13480.66 (± SD) reads per sample. ASVs were generated and subsequently taxonomically classified into genera (Fig. 1A). Despite regional differences, several core genera were consistently present in flies from all locations. The genus *Dysgonomonas* was the most prevalent, being detected in all samples, while *Vagococcus*, was the most abundant feature, with a prevalence of 98.44%, followed by *Pseudomonas* (91.47%), *Ignatzschineria* (82.17%), *Providencia* (79.84%), and *Lactobacillus* (77.51%). Chungnam presented the highest average relative abundances of *Dysgonomonas* and *Vagococcus* (Fig. 1B). Interestingly, *Dysgonomonas* average relative abundance was the lowest in Chungbuk while *Vagococcus* was the lowest in Jeonnam. *Pseudomonas* was highly prevalent in Seoul. Similarly, Seoul, Incheon and Gyeonggi exhibited a marked abundance of *Ignatzschineria*. *Providencia* showed the highest abundance in Chungbuk and Jeonnam, whereas *Enterococcus* showed high abundance in Chungbuk and Chungnam. Furthermore, *Proteus* was particularly abundant in Seoul and Gyeonggi. Relative abundances of the top 100 most abundant ASVs are presented in (supplementary Table S2).

**Figure 1.**
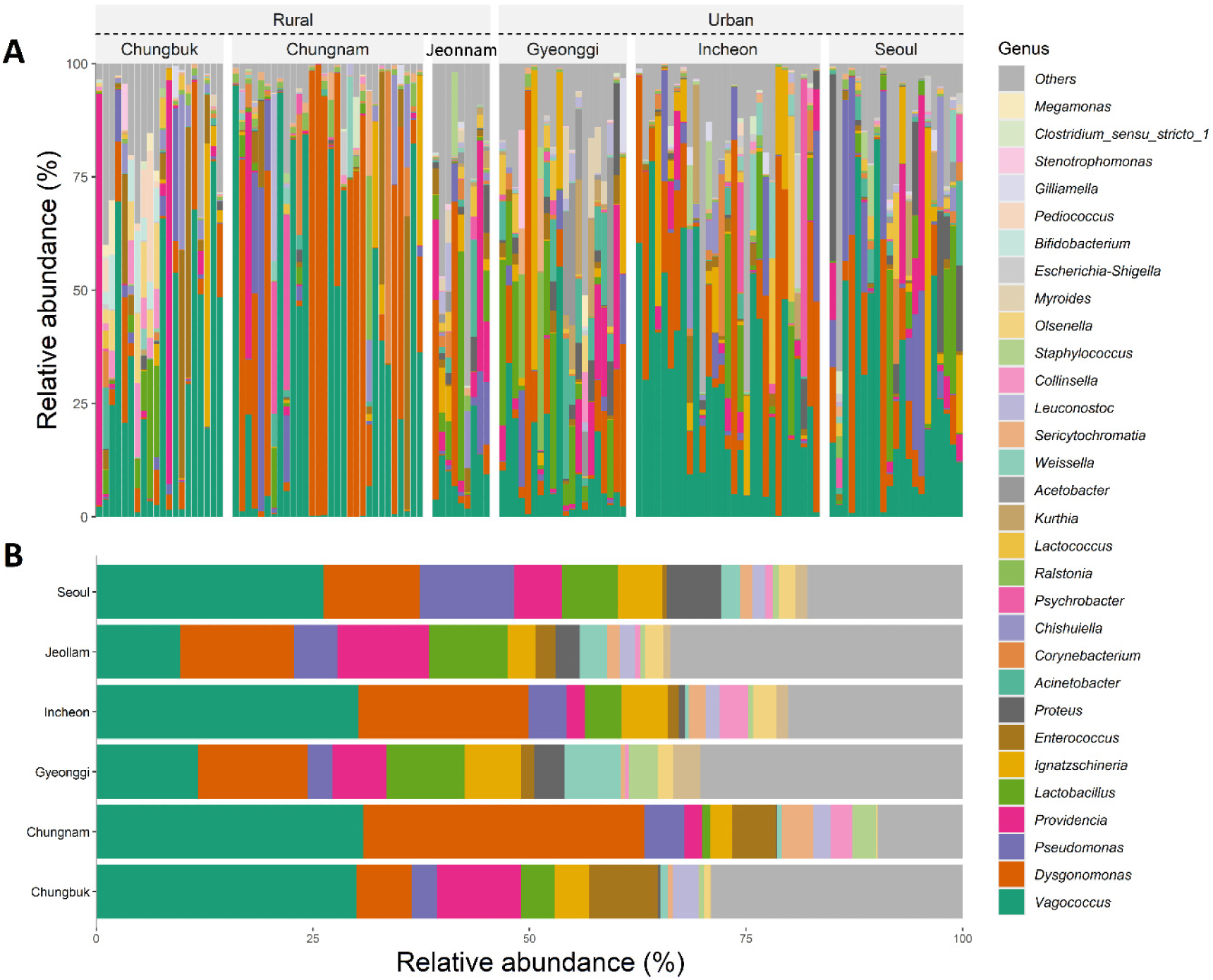
Microbiome composition of *Lucilia sericata* from six regions in South Korea. Relative abundances of the top 30 most abundant bacterial ASVs found in flies from rural and urban locations (A). Average ASV relative abundances by province of sampling showing the top 15 most abundant bacterial ASVs found in *L. sericata* (B).

### Core microbiome

A total of 777 (35.4%) ASVs were unique from urban locations. Rural areas had 851 (38.8%) unique ASVs, while they both shared 564 (25.7%) features (Fig. 2A). Similarly, the core microbiome across all provinces was composed of 92 (4.2%) (Fig. 2B). Chungnam had 356 unique ASVs representing 16.2% of the total features, followed by Gyeonggi (12.6%), Seoul (11.8%), Jeonnam (9.9%), Chungbuk (9.7%) and Incheon (8.1%). Overall, the shared features between individual provinces did not surpass 50 ASVs.

**Figure 2.**
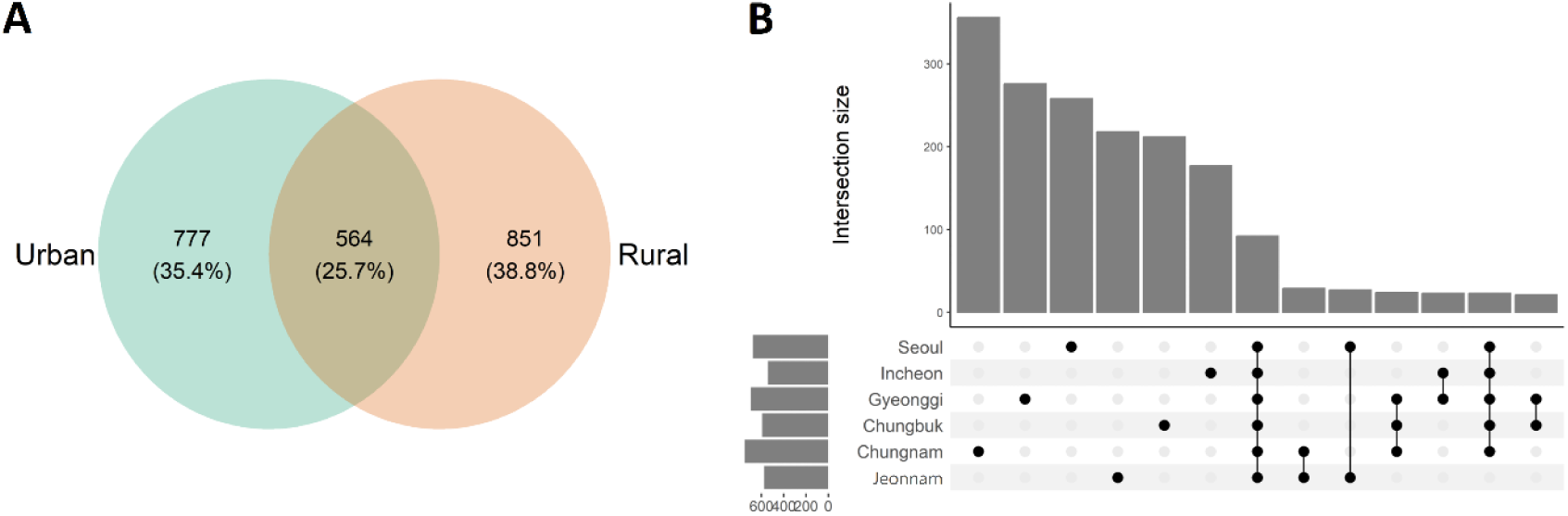
Shared bacterial ASVs from the microbiome of *Lucilia sericata* sampled in South Korea. Venn diagram showing the number of shared and unique ASVs in flies sampled from urban and rural locations in South Korea (A). Upset plot of shared and unique ASVs found in flies classified by sampling region (B).

#### Alpha Diversity of the bacterial communities

ASV richness was lower in Incheon compared to Seoul (*p* = 0.031), Gyeonggi (*p* = 0.015) and Jeonnam (*p* = 0.006) (Fig. 3A). Richness in Jeonnam samples was higher than in Chungnam (*p* = 0.027), while it was higher in Gyeonggi than in Incheon (*p* = 0.013). There were no significant differences between Gyeonggi, Jeonnam, Chungbuk and Seoul. Shannon index was significantly lower in Chungnam compared to Chungbuk (*p* = 0.015), Incheon (*p* = 0.013), Seoul (*p* = 0.006), Gyeonggi (*p* = 0.006), and Jeonnam (*p* = 0.006) (Fig 3B).

**Figure 3.**
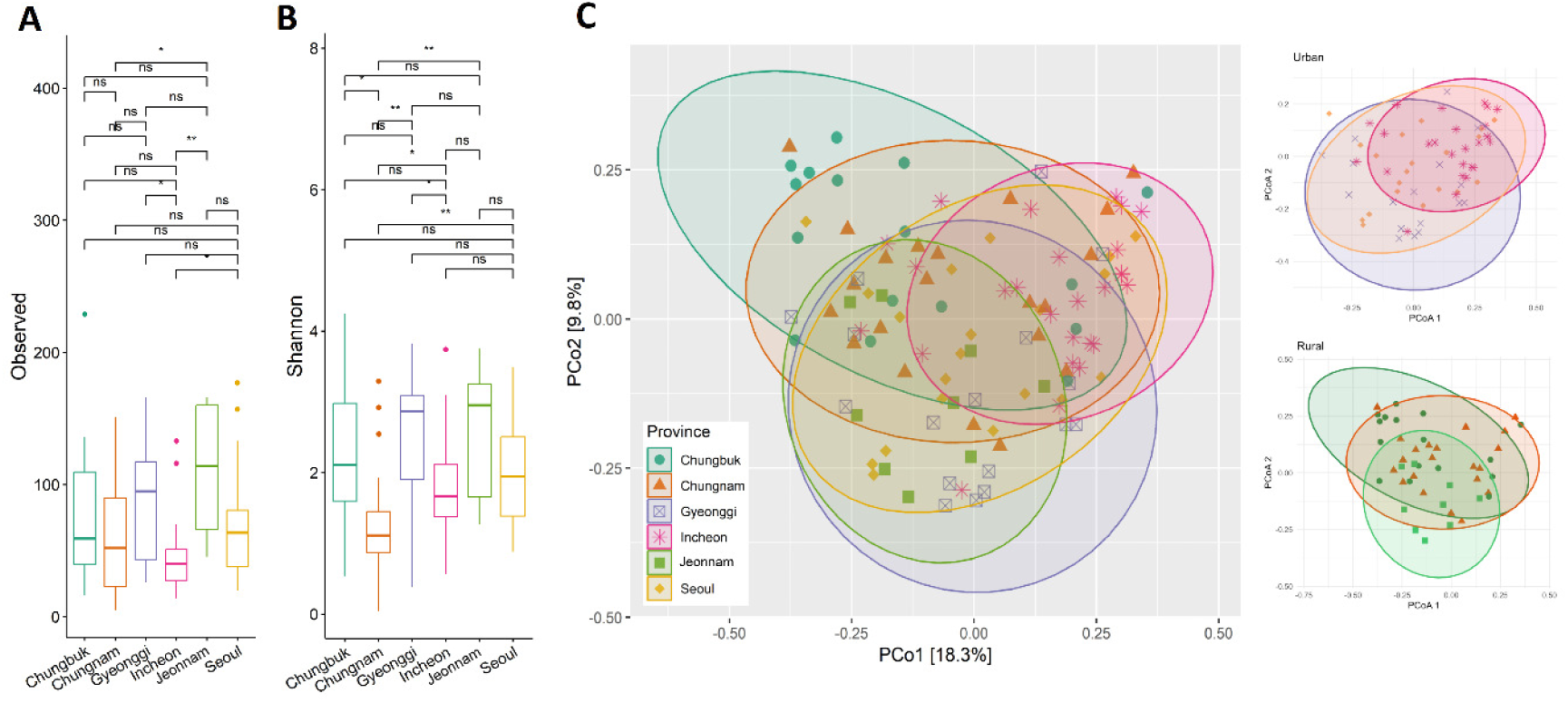
Diversity comparisons of the microbiome in *Lucilia sericata* flies. Fly microbiome richness compared among flies collected across sampling region (A). Fl microbiome Shannon index compared among flies collected across different regions (B). Principal component analysis (PCoA) of the unweighted UniFrac distances of the microbiome of flies collected across different regions, panels with urban and rural samples plotted separately are shown on the right side for better visualization (C).

#### Beta Diversity of the bacterial communities

PERMANOVA of the weighted UniFrac distances revealed differences in microbiome composition by province (*p* = 0.001) (Fig. 3C, supplementary Figure S2). Pairwise PERMANOVA detected significant differences (*p* < 0.05) among all provinces except for Gyeonggi and Seoul (*p* = 0.05) (*p* = 0.175) (supplementary Table S3). The LEFSE analysis was used to identify enriched taxa found in each province (Fig. 4). Seoul was characterized by *Paraclostridum*, *Comamonas* and *Selenomonas*. Simmilarly, Incheon had *Gluconobacter* as the only enriched genus. Chungnam was marked by *Ralstonia*, *Microbacterium* and *Bradyrhizobium*. The taxa with the highest LDA in Gyeonggi were *Acinetobacter*, *Proteus* and *Kurthia*, in Jeonnam were *Pseudomonas*, *Providencia* and *Lactobacillus*, while in Chungbuk were *Olsenella*, *Bifidobacterium* and *Collinsella*.

**Figure 4.**
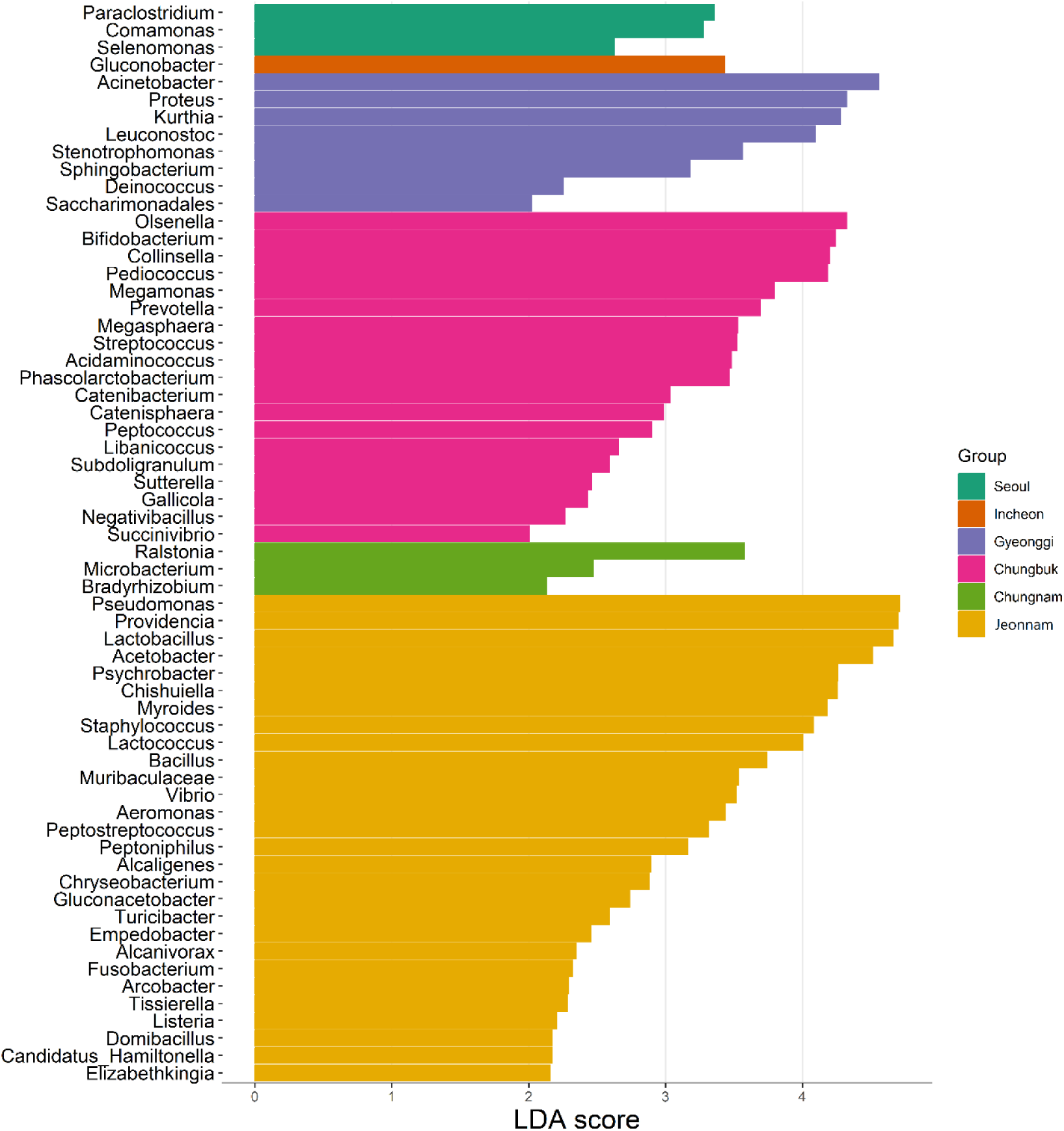
Linear discriminant analysis (LDA) effect size plot of the enriched taxa in the microbiome of *Lucilia sericata* flies sampled in South Korea, compared by regions. Enriched taxa with LDA > 2 and *p* < 0.05 are shown.

### Detection of Proteus mirabilis and Providencia spp. in flies

Metabarcoding revealed a high prevalence of *Proteus and Providencia* ASVs. *Proteus mirabilis* was detected in 76 of 129 flies, representing a prevalence of 100% in Jeonnam, 80% in seoul, 70% in Gyeonggi, 55% in Chungbuk, 48% in Incheon and 36% in Chungnam. Similarly, *Providencia* was detected in 103 samples, with a prevalence of 100% in Jeonnam, 85% in Seoul, 85% in Chungbuk, 82% in Incheon, 75% in Gyeonggi, and 36% in Chungnam. Subsequent PCR screening with species-specific primers confirmed *Proteus mirabilis* in the samples*. Providencia* amplicons using genus-specific primers were used to construct a phylogenetic tree in which samples grouped with *P. rettgeri/P. vermicola, P. huaxiensis, P. rustigianii/P. alcalifaciens*, *P. manganoxydans*, and *P. stuartii.* (supplementary Figure S3).

### Antimicrobial resistance genes profile and correlation with bacterial community

Among the ten tested genes, *tetA* and *vanA* were not detected in any sample (Fig. 5A). The ARGs and integron genes profile are shown in Table 1. PERMANOVA revealed significant differences by group (*p* = 0.035). Differences of ARGs and integron composition were found between Incheon with Gyeonggi (p = 0.018), Incheon with Jeonnam (p = 0.021) and Gyeonggi with Chungbuk (p = 0.004) The network correlation analysis (Fig. 5B) revealed complex interactions between bacterial taxa and antimicrobial resistance genes, encompassing both positive and negative associations of varying significance. *Lactobacillus* and *Enterobacteriaceae* showed a significant positive correlation with *intI*. Several taxa were associated with *intII*, including *Leucobacter*, *Lactococcus*, *Acinetobacter*, and *Proteus*. Four ARGs exhibited notable positive correlations with specific bacterial groups *ermB* was positively associated with *Providencia*, *Enterobacterales*, and *Wohlfahrtiimonadaceae*, while *sul1* correlated positively with *Morganellaceae* and *Sphingobacterium*. The *aac(6′)-Ib-cr* gene showed positive correlations with multiple taxa, including *Proteus* and *Myroides*. Similarly, *cat1* was positively associated with *Leucobacter*, *Lactococcus*, *Acinetobacter*, and *Proteus* (supplementary Figure S4). In contrast, *mecA* was negatively correlated with *Ignatzschineria* and *Proteus*. However, these associations were not statistically significant (p = 0.059 and p = 0.062, respectively). Negative co-occurrence relations are shown in supplementary Figure S5.

**Figure 5.**
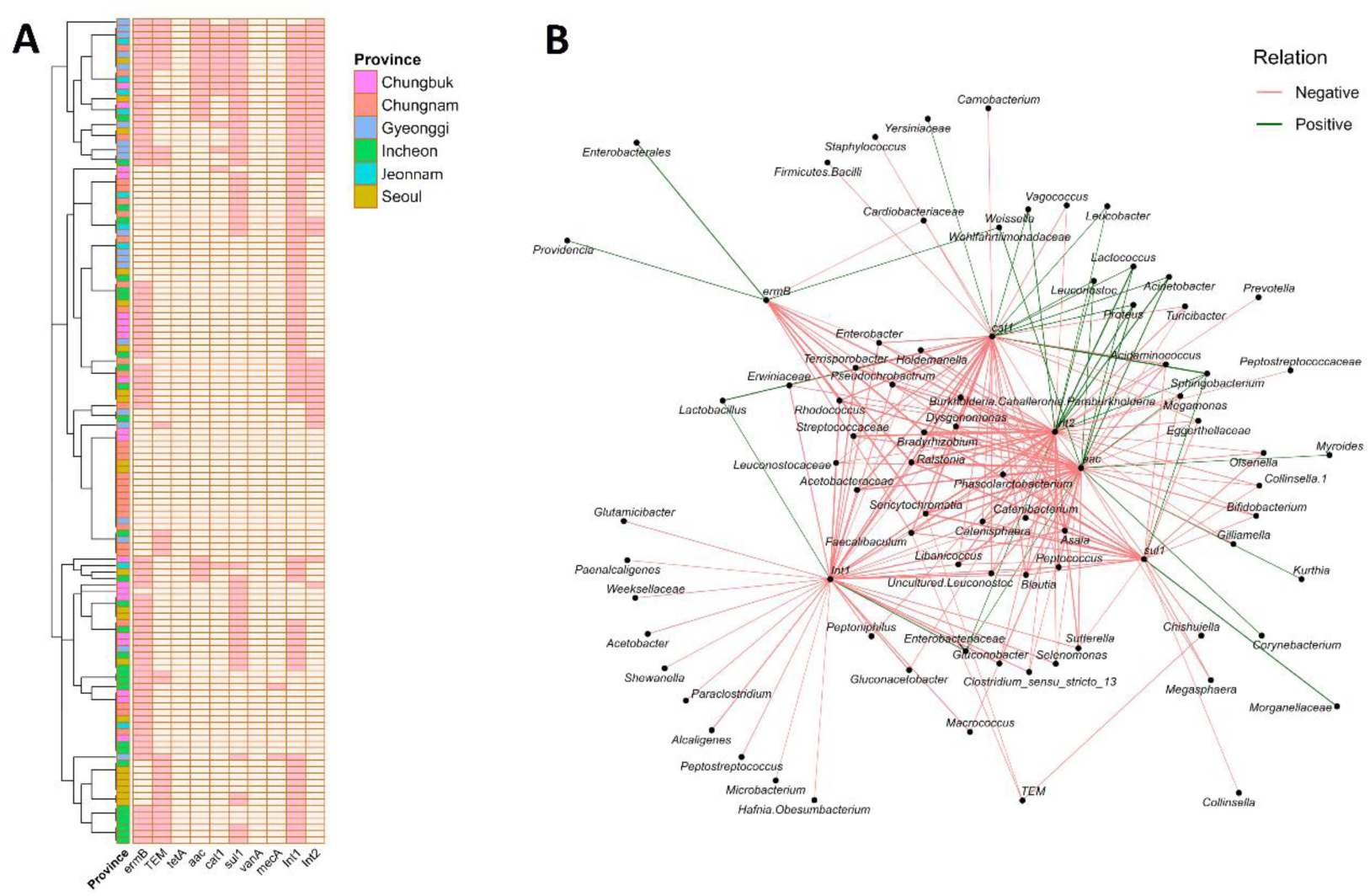
Resistome profile of the *Lucilia sericata* microbiome in South Korea. Clustered heatmap of presence (dark pink) and absence (light pink) of antibiotic resistance genes (ARGs) across samples from six regions in South Korea, rows (samples) are hierarchically clustered based on Euclidean distance and complete linkage grouping samples with similar ARG profiles, columns (genes) are not clustered, row colors indicate sampling location (A). Spearman correlation network of ARG presence (binary) and ASV abundance across *Lucilia sericata* samples from South Korea, edges represent significant correlations where *p* < 0.05 and absolute correlation coefficient |ρ| > 0.2, edge thickness is proportional to correlation strength (B). The *aac* legend refers to the *aac(6′)-Ib-cr gene*.

## Discussion

Flies are hosts to diverse microbial communities reflecting their interactions with various environments. While human pathogens have consistently been identified in house flies (*Musca domestica*) (19), other flies such as ***Lucilia sericata*** are not extensively studied. Due to their necrophagous lifestyle this genus is exposed to unique bacterial communities, making them likely carriers of pathogenic and opportunistic bacteria (11). Here, we assessed the *L. sericata* microbiome from locations ranging from urban centers (Seoul, Incheon) to semi-urban (Gyeonggi) and rural (Chungbuk, Chungnam and Jeonnam) areas.

Core genera including *Dysgonomonas, Vagococcus, Pseudomonas, Ignatzschineria, Providencia*, and *Lactobacillus* were detected across all provinces, indicating their potential symbiotic role within the *L. sericata* gut (20). *Dysgonomonas*, a genus commonly linked to anaerobic fermentation and decomposition processes (21), was detected across all regions, with the highest relative abundance observed in Chungnam and the lowest in Chungbuk. Similarly, *Vagococcus,* the most abundant genus overall, also reached its highest relative abundance in Chungnam. As a lactic acid bacterium frequently isolated from decomposing matter (22), its co-occurrence with *Dysgonomonas* may suggest potential indirect symbiotic interactions. Jeonnam, however, showed the lowest abundance of *Vagococcus*, but presented high abundances of other lactic acid bacteria and the most diverse microbiome in the dataset. *Pseudomonas* often linked to anthropogenic and aquatic environments (23) was especially enriched in Seoul, Dangsan-dong which is located near a water body, and Yeonseo Market, a busy area with high human activity. These conditions create a suitable environment for environmental *Pseudomonas* to be acquired by flies. Similarly, **the** bacteremia**-producing emergent** pathogen *Ignatzschineria* (24) was abundant in the urban regions of Seoul, Incheon, and Gyeonggi. Several of other dominant genera identified in the present study are consistent with those reported in previous investigations of blow fly microbiota. *Lactobacillus*, was found in a previous study at high abundance in the salivary glands of *Lucilia sericata* (11). It can inhibit other pathogens through acidification and direct antimicrobial peptide production (25). Similarly, **the antimicrobial peptide-producing** *Myroides* (26) ha**s** been identified in the bacterial community of *L. sericata* (11) and other fly species (26–28).

Significant geographic variations in bacterial alpha diversity were found across regions. Chungnam exhibited lower diversity compared to all other provinces, suggesting a more uneven microbial distribution likely influenced by *Vagococcus* and *Dysgonomonas* dominating the microbiome across samples. Gyeonggi and Jeonnam had higher bacterial richness, while Incheon was consistently less diverse. Jeonnam, is characterized by rural and agricultural environments, and Gyeonggi, with its mix of urban and peri-urban areas, likely promoting richness. While both Gyeonggi and Jeonnam exhibited high richness, no significant differences were found between them, Chungbuk, and Seoul. Interestingly, in the case of Jeonnam, samples were collected in the island of Jindo, providing a source of a more distinct set of environmental microbes. Beta diversity analysis demonstrated statistically significant differences by province. The absence of significant differences between Gyeonggi and Seoul (PERMANOVA, *p* = 0.175) may be attributed to their geographic proximity and shared urban features, potentially leading to comparable microbial exposures and ecological conditions.

Differential abundance analysis identified region-specific bacterial signatures. Jeonnam flies harbored the highest number of enriched taxa. Previous studies indicate that flies from agricultural environments exhibit higher microbial diversity (29). In contrast, the microbiome of flies from urban, Semi-urban areas such as Seoul, Incheon and Gyeonggi were characterized by genera including *Comamonas*, *Gluconobacter* and *Acinetobacter*, which are frequently associated with human-influenced environments, food and waste (30–32). *Acinetobacter* **is** an environmental bacterium **that can** colonize the gut of ticks after antibiotic-induced dysbiosis (33) and **is known to** elicit antimicrobial responses in *L. sericata* (34). Analysis of the core microbiome revealed that only 4.2% of amplicon sequence variants (ASVs) were shared across all provinces. Comparison by land use type indicated a higher proportion of unique ASVs in rural (38.8%) versus urban (35.4%) locations, with only 25.7% shared, further highlighting the influence of local habitat on microbial community structure. Although Chungnam exhibited the lowest microbial richness among all regions, it surprisingly harbored the highest number of unique ASVs (16.2%). Diverse environmental factors such as food sources, pollution, or antimicrobial exposure may favor the colonization of different microbes from different environments. Overall, changes in microbiome diversity across regions and the small core microbiome found across all samples may reflect the capacity of blow flies to acquire and disseminate microbes from their environment (20).

Clinically relevant taxa, such as *Proteus mirabilis, Providencia, Helicobacter, Clostridium, Vibrio,* and *Myroiodes,* some of which are causative agents of nosocomial, gastrointestinal and urinary tract infections (26, 35), were abundant in the fly microbiome. *Proteus mirabilis* and *Providencia stuartii* are two bacteria that are clinically relevant as they are commonly found in blow flies and are known to produce coinfections in humans (36–38). *Providencia* was most abundant in Chungbuk and Jeonnam, both of which are rural regions. Notably, *Providencia* is known to produce multiple xylanases, facilitating the breakdown of xylan, a compound frequently present at decomposition sites (39), pointing at important physiological symbiotic roles in their hosts. *Proteus* was highly abundant in Seoul and Gyeonggi. *P. mirabilis* exhibits swarming behavior and produces a distinct odor that attracts blowflies (40). It has been previously identified as a dominant bacteria associated with *Lucilia sericata* and *Lucilia cuprina* (11). According to a previous studies *Providencia* and *Vagococcus* are the most dominant genera across all life stages in *L. sericata* (37). *Proteus mirabilis* is an important causative agent of urinary tract infections (41) and is responsible for the majority of infections caused by *Proteus* species (42). It is also a known cause of bacteremia and other hospital-acquired infections (43). Notably, associations between *P. mirabilis* and autoimmune conditions such as rheumatoid arthritis have also been reported (44). The management of *P. mirabilis* infections is increasingly challenged by the emergence of extended spectrum beta-lactamase producing strains (45). Both resistant and sensitive *P. mirabilis* strains are able to persist in the fly guts suggesting the digestive tract environment serves as reservoirs for ARG carrying *P. mirabilis* (46). Similarly, *Providencia* has been identified as a notable opportunistic pathogen responsible for a broad range of hospital acquired infection (43). It has been reported as the fourth most common gram-negative pathogen in UTIs, commonly displaying antimicrobial resistance (47). A number of these species, such as *P**rovidencia** stuartii* and *P**rovidencia** rettgeri*, are known for causing hospital-acquired infections and for their resistance to various antibiotics (48).

Other pathogenically relevant taxa detected here, such as *Helicobacter*, have previously been associated with *L. sericata* (3), whereas other such as *Vibrio* species have not been commonly reported in association with blow flies. In this study, *Vibrio* was detected in 14 out of 129 fly samples, with the highest read counts originating from the fish market area in Jeonnam. Since *Vibrio* species are commonly associated with marine environments and seafood waste (49), their presence could be related to the exposure to caught fish implying that spatial factors contribute to variations in the bacterial profiles of the flies.

The *ermB* gene was the most frequently detected ARG across all regions, with highest prevalence in Incheon (79.3%) and Chungbuk (75%). It has been detected before in houseflies (6) as well as in livestock (50). *TEM* showed high overall prevalence in urban areas compared to rural areas while *aac(6′)-Ib-cr* showed higher detection in Jeonnam (55.5%). *Sul1* showed comparable presence in all samples indicating that sulfonamide resistance is broadly disseminated in both urban and rural environments. In rural and agricultural areas of South Korea, the use of manure is still common and can contribute to the environmental spread of antibiotic resistance genes (51). Interestingly, *aac(6′)-Ib-cr* and *cat1* exhibited high prevalence within Jeonnam. These patterns may reflect region-specific persistence of these resistance genes. In contrast, the *tetA* and *vanA* genes were not detected in any sample, suggesting limited transmission of tetracycline and vancomycin resistance. This aligns with previous findings indicating that resistance genes to tetracycline and vancomycin genes are rarely associated with integrons (18). Moreover, vancomycin is most commonly used in hospital settings (52). The low detection rate of *mecA* may suggest a limited environmental reservoir of methicillin-resistant *Staphylococcus aureus* (MRSA) and related bacteria outside clinical settings. However, two positive samples were detected in this study. Additionally, *L. sericata* produces antimicrobial peptides protective against MRSA and methicillin-sensitive *Staphylococcus aureus* (MSSA) (53) as well as vancomycin-resistant *Enterococcus* (54). The class 1 integron (*intI*) was detected at high frequencies in all regions, especially in Jeonnam (88.9%) and Seoul (76.2%). It is a key driver of ARGs dissemination in environmental bacterial communities and is commonly found in plasmids (55). Among the multiple antibiotic resistance genes tested, *cat1* was consistently detected in samples where class 1 integrons were present. Class 2 integrons (*intII*) were also present, with Gyeonggi (60%) and Jeonnam (55.5%) exhibiting particularly high prevalence. Class 2 integrons are related to a more conserved set of ARGs such as *Acinetobacter*, *Enterobacteriaceae*, *Salmonella* and *Psuedomonas* and associated with the Tn7 transposon family (56). Notably, class 2 integrons are found in *Proteus* (56). Further, correlations between *Enterobacteriaceae* and both integrons are consistent with previous findings indicating that *Enterobacteriaceae* isolated from sewage can serve as reservoirs of integron associated antibiotic resistance genes (57). Moreover, *ermB*, *sul1*, *aac(6′)-Ib-cr*, and *cat1* were positively associated with specific taxa that exhibit multi-drug resistance such as *Proteus*, *Providencia*, *Enterobacterales*, *Morganellaceae*, and *Myroides*, suggesting these bacteria may contribute to ARG maintenance and transmission (26, 58, 59).

Limitations of this study include compositional bias and small sample sizes that can potentially mask relevant associations. Co-relation networks are not causal and therefore can only infer relations based on co-occurrence patterns. Additionaly, while PCR-based detection or ARGs allows for the identification of genes and integrons, it does not assess actual phenotypic resistance. In contrast, traditional culturing methods can test functional resistance but are limited in depth and often fail to capture fastidious taxa that are detected by metagenomics.

Here, we showed that *L. sericata* harbors a diverse microbiome and resistome shaped by regional biotic or abiotic factors. The presence of pathogenic taxa and ARGs across both urban and rural regions highlights their ecological plasticity and their capacity to serve as reservoirs. Our findings add to the growing body of evidence that blow flies may disseminate antibiotic resistance in their microbiome.

## Acknowledgements

This research was supported by the National Research Foundation of Korea (NRF) grant funded by the Korea government (MSIT) (No. RS-2024-00456300) and by a grant of the Korea Health Technology R&D Project through the Korea Health Industry Development Institute (KHIDI), funded by the Ministry of Health & Welfare, Republic of Korea (grant number: RS-2024-00406488).

## Author contributions

Conceptualization: AS, XC, and JYK. Methodology: AS, XC, JHC, MY, and JYK. Investigation: AS, XC, JHC, SO, MK, DK, DYC, and YHC. Formal analysis: AS, and XC. Writing – original draft: AS, and XC. Writing – Review & Editing: XC, and JYK. Supervision, project administration, and Funding acquisition: KJY.

## Data availability

Raw sequencing data and associated metadata are available in the NCBI Short Read Archive under the BioProject number PRJNA1274159.

